# Subtype-specific autophagy profiles in renal cell carcinoma

**DOI:** 10.64898/2026.11.17.767069

**Authors:** Feyza Kasikci, Julie Aarmo Johannessen, Eva Bassols-Citores, Maria Frison, Siri Andresen, Elise Komttebed, Frode S. Nilsen, Stig Müller, Karol Axcrona, Leiv Rønneberg, Manuela Zucknick, Lorant Farkas, Jorrit M. Enserink, Helene Knævelsrud

## Abstract

Autophagy is a highly conserved cellular mechanism essential for recycling intracellular components, providing energy and maintaining cellular homeostasis. The role of autophagy in cancer is double sided. It can suppress tumorigenesis by degrading damaged organelles and preventing chronic inflammation, while enabling survival of existing tumor cells during starvation and stress, and thus support tumor growth. The role of autophagy in renal cell carcinoma (RCC), the most common type of kidney cancer, remains insufficiently described. RCC originates from epithelial cells of the renal tubule and is divided into histological subtypes where clear cell RCC is the most common, followed by papillary and chromophobe RCC. Recent studies of RCC suggest that mTORC1 and autophagy are involved in kidney cancer development and response to treatment, advocating the need to better understand the role of autophagy in RCC. To this end, we investigated autophagy markers by immunostaining, immunoblotting and quantitative PCR in biobanked fresh-frozen tissue samples of RCC tumor and matched normal tissue. Overall, we found reduced levels of LC3B and GABARAPL1 in clear cell RCC, in stark contrast to chromophobe RCC and benign oncocytoma. This suggests that autophagy levels differ between renal tumor subtypes, indicating that autophagy could play subtype-specific roles. In addition, autophagy modulating drug assays demonstrated autophagy dependencies in metastatic RCC cell line spheroids. This could provide new inroads for therapeutical approaches in RCC.

## Introduction

Renal cell carcinoma (RCC) arises from the tubular lining of the nephrons in the kidneys and is the main type of kidney cancer (Bahadoram & Davoodi, 2022). RCC is more common in men and has a mortality rate of around 30-40% with more than 200 000 new cases and increasing worldwide each year (Siegel et al., 2019). Most renal tumors are discovered by chance by medical imaging for other indications (Padala et al., 2020). Some of these will be oncocytomas, a benign solid renal tumor, but most are malignant RCC. The World Health Organization (WHO) has classified RCC in different subtypes based on their histological appearance (Humphrey et al., 2016). The most common of these are called clear cell, chromophobe and papillary RCC. Clear cell RCC has the highest frequency with 70% of all kidney malignancies and also has the worst prognosis (Bahadoram & Davoodi, 2022). Primary RCC is treated by surgery and metastatic RCC by combinations of immunotherapy and targeted therapies (Powles et al., 2024). These include tyrosine kinase inhibitors, immune check point inhibitors and mTOR inhibitors. Although these treatments significantly prolong survival, most patients eventually progress. Cancer cells have been shown to exploit autophagy to withstand the cytotoxicity of cancer drugs, suggesting that autophagy inhibition could potentially help overcome drug resistance in RCC (Grimaldi et al., 2015).

Autophagy is a conserved cellular mechanism that has a crucial role in cellular homeostasis (Nakatogawa, 2020). Autophagy has a dual role in cancer, both acting as a tumor suppressor, since absence or defect in autophagy causes accumulation of cellular damage and genomic instability (White, 2015), and as a survival mechanism which can promote tumorigenesis depending on tumor stage, cancer type and mutations (Shin, 2020). Upon autophagy initiation, a portion of the cytosol will be engulfed by the forming autophagosome. The autophagosome subsequently fuses with a lysosome, whereby the cargo is degraded and released into the cytoplasm for recycling (Yamamato et al., 2023). Autophagosome formation is marked by the lipidation of cytosolic Atg8 family proteins. In mammalian cells these are Microtubule associated protein 1 light chain 3 (MAP1LC3) alpha (MAP1LC3A/LC3A), MAP1LC3B/LC3B, MAP1LC3C/LC3C, GABA-type A receptor-associated protein (GABARAP), GABARAP like 1 (GABARAPL1) and GABARAPL2 (Lamb et al., 2013). Atg8 proteins are bound by cargo-receptor proteins, such as sequestosome 1 (SQSTM1/p62), which also recognize and bind autophagy substrates, enabling their degradation by autophagy. SQSTM1/p62 can also recruit the ULK1 kinase complex to promote autophagosome formation (Zaffagnini & Martens, 2016) and SQSTM1/p62 is itself turned over by autophagy.

Autophagy is regulated by a complex network of signaling pathways that respond to the cell’s nutrient availability, energy levels, stress status, and growth (Shpilka et al., 2011). A key complex for the initiation of autophagy is the ULK1 kinase complex (He & Klionsky, 2009). Autophagy is also negatively regulated by mTORC1 (mechanistic target of rapamycin complex 1). Under nutrient-rich conditions mTORC1 phosphorylates targets including ULK1, S6K and 4E-BP1 to inhibit catabolism and promote anabolism. mTORC1 is positively activated by PI3K/AKT pathway (He & Klionsky, 2009). There are currently a few drugs that affect autophagy and are used in the clinic, but not yet in cancer treatment. Chloroquine (CQ) and hydroxychloroquine (HCQ) are two FDA approved drugs that inhibit autophagy, although they have also many other effects (Shin, 2020). CQ and HCQ inhibits the fusion of autophagosomes with lysosomes (Mauthe et al., 2018). Traditionally, HCQ is used to treat malaria and inflammatory diseases (Daily et al., 2022), however its anti-tumor effects, particularly when combined with other anti-cancer drugs such as those for breast (Cave et al., 2018) and colon (Liu et al., 2019) cancer, have highlighted its potential in future cancer therapies. Although these drugs are not yet approved for cancer treatment, several clinical trials are underway (Mohsen et al., 2022). For instance, a phase I/II clinical trial observed tolerance and disease control with HCQ in combination with mTORC1 inhibitor everolimus in advanced RCC (Haas et al., 2019).

A few studies have investigated autophagy in RCC cell lines. Some of these conclude that RCC cell lines exhibit elevated autophagy levels (Bray et al., 2012; Guo et al., 2019). However, another study rather implies autophagy as a suppressor of RCC development (Wang et al., 2018). Autophagy in RCC patient samples remains poorly characterized, especially in non-clear cell RCC subtypes. We hypothesized that RCC subtypes display differential autophagy levels and dependency.

## Results

### Characterization of patient samples included in this study

To investigate autophagy markers in different subtypes of RCC, we acquired representative samples from a biobank of fresh-frozen kidney tumor samples with corresponding normal kidney tissue. Each tumor sample was inspected by a pathologist to ensure that they were representative of their histological classification, devoid of necrosis and contained almost exclusively tumor tissue (representative images in Figure 1A). Clear cell RCC is characterized by lipid-rich cytoplasm, which when washed away during staining, renders the cytoplasma clear in appearance. Chromophobe RCC is identified as large pale cells typically with wrinkled hyperchromatic nuclei, reticulated cytoplasm and perinuclear halos. Papillary RCC has a typical morphology of papillary or tubular pattern with basal membrane surrounded by a single layer of cuboidal cells with scant or lightly basophilic cytoplasm. Oncocytomas are known to be more heterogenous, however densely granular eosinophilic cytoplasm and tubular structures are common. We selected samples from three different subtypes of RCC: Clear cell (10 samples), papillary (11 samples) and chromophobe (7 samples) and 10 samples of oncocytomas, a benign form of kidney tumor (Fig. 1B).

**Figure 1:**
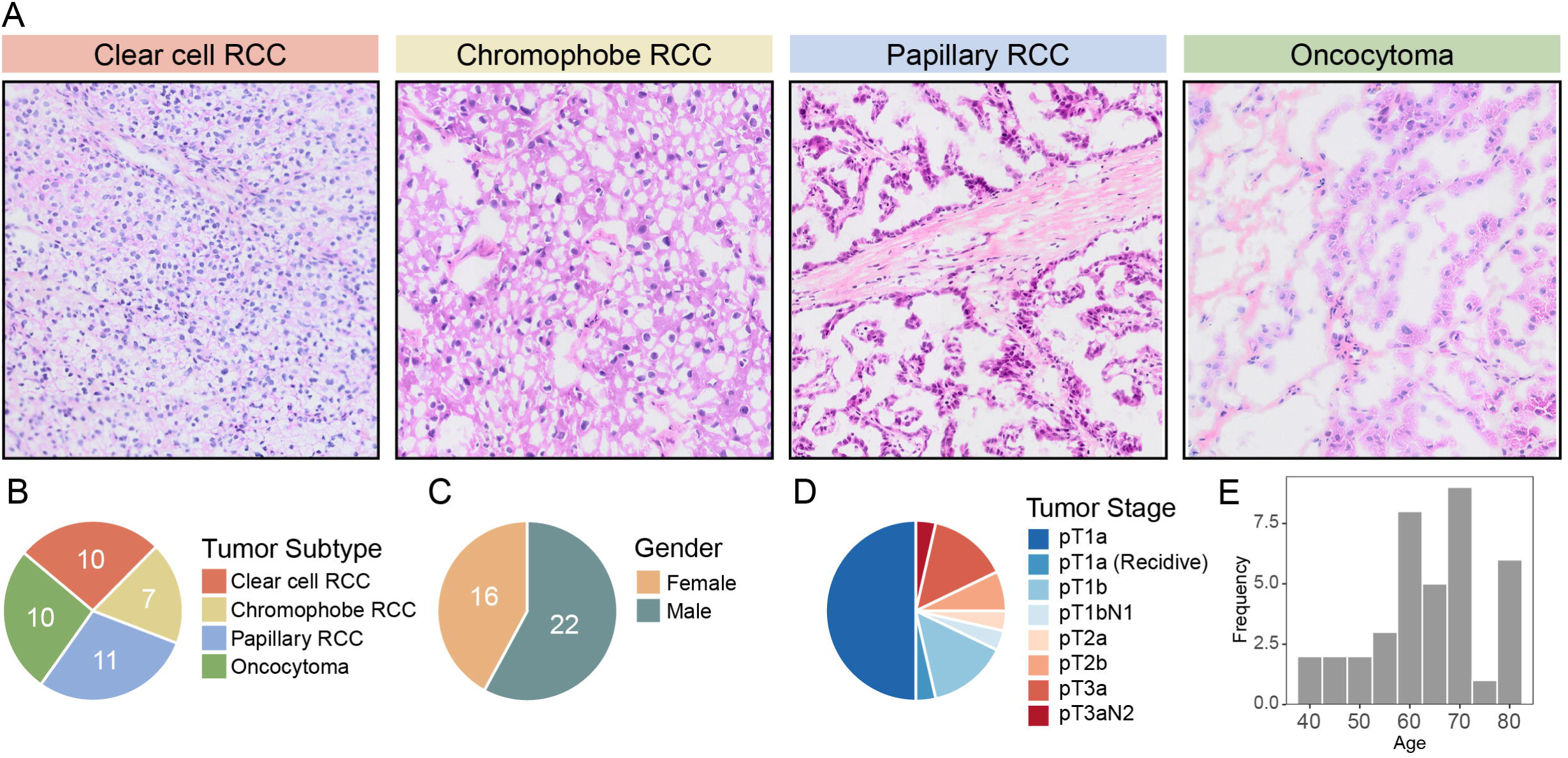
Overview of biobanked samples included in this study. **A**: Representative images of HPS-stained sections of different subtypes of renal tumors represented in the biobank. Each tumor sample has a corresponding normal kidney sample (not shown) from the same patient. **B-D**: Distribution of kidney tumor subtypes, gender and tumor stages shown as pie charts. **E**: Age distribution shown as histogram.

From the 38 cases used in this study, 22 patients were male and 16 females (Fig. 1C). Most of the tumors (around 70%) were in stage 1, with a few in stage 3 (around 20%) and even fewer in stage 2 (around 10%, Fig. 1D). The age distribution of the patients was centered around 60-80 years, with some in the 40–50-year bracket (Fig. 1E). The distribution of age, gender and stage of our samples are representative of the epidemiology of RCC in general: the majority of RCC patients are male and above 60 years of age, and most tumors are caught in earlier stages (Bahadoram & Davoodi, 2022). These biobanked samples were investigated by different methods to assess the involvement of autophagy (Fig. S1A).

### Renal tumor subtypes show differences in autophagy markers on immunostaining

To assess levels of autophagy associated markers, we stained our cryo-sectioned patient samples with antibodies against LC3 and p62 (Fig. 2A-D and Fig. S2A). Despite being raised against human LC3B, we cannot exclude detection of LC3A and LC3C with this antibody and therefore refer to this as LC3. We found that LC3 and p62 staining was more prominent in normal kidney tissue than in tumor tissue in clear cell RCC (Fig. 2A). In contrast, chromophobe RCC generally displayed the opposite pattern, with higher levels of LC3 and p62 in tumor compared to its normal kidney counterparts (Fig. 2B). Papillary RCC samples had slightly higher levels of LC3 and more abundant p62 in tumor than normal tissue, similar to chromophobe RCC samples (Fig. 2C). Oncocytoma displayed more heterogenous staining patterns, although higher levels of p62 were generally observed in tumor tissue (Fig. 2D). Quantification of LC3 intensity between tumor and normal tissue for the subtypes revealed no statistically significant differences among the subtypes (Fig. 2E), whereas p62 levels in clear cell RCC significantly differed from the other subtypes, where clear cell RCC consistently showed lower p62 levels in tumor compared to normal tissue (Fig. 2F). Overall, we found that the pattern of these autophagy proteins as detected by immunofluorescence more pronouncedly differed by RCC subtype for p62 than LC3.

**Figure 2:**
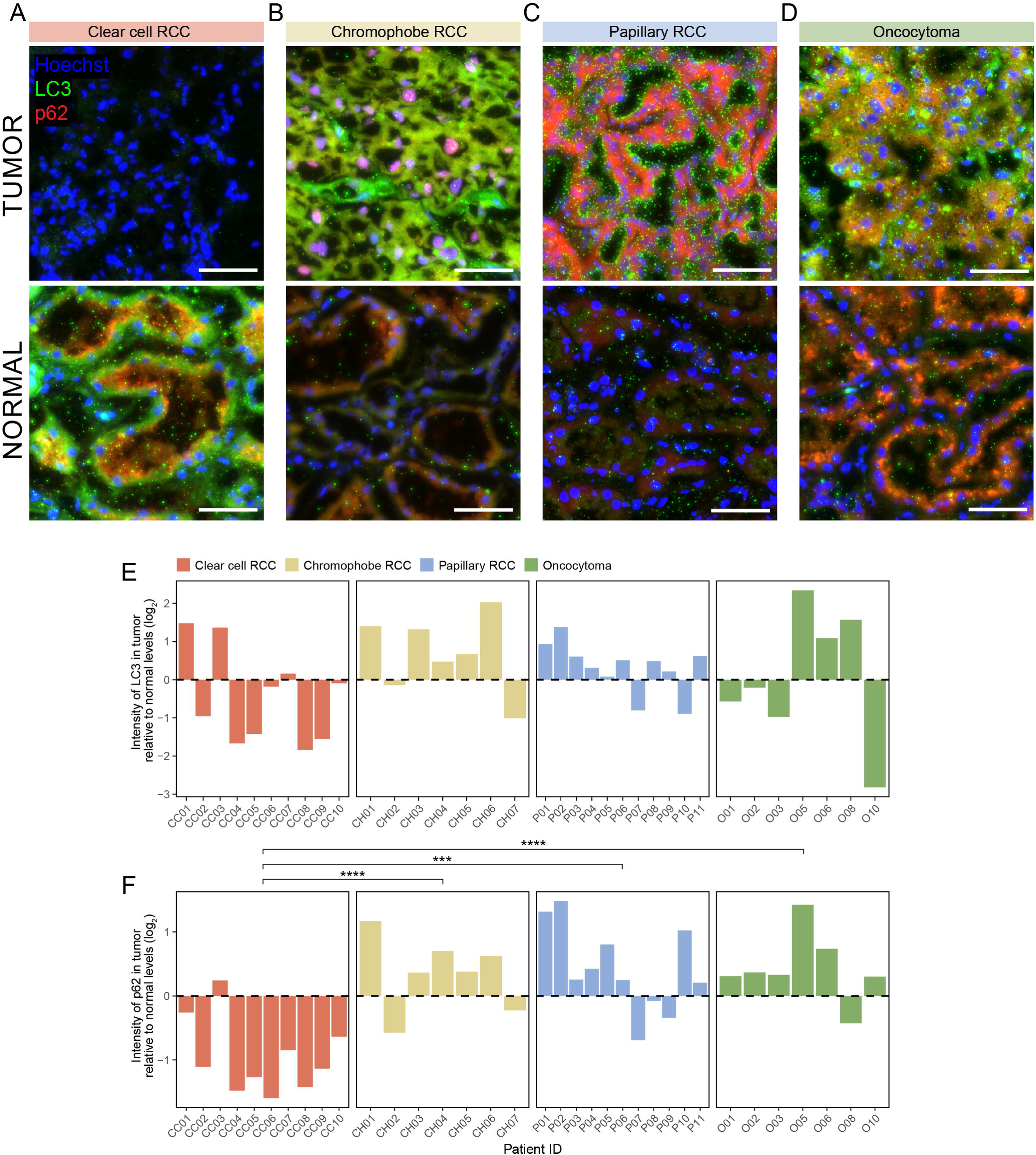
Immunofluorescence shows decreased LC3 and p62 levels in clear cell RCC. **A-D:** Representative immunofluorescence images of tumor tissue from kidney tumor subtypes and adjacent normal tissue stained for autophagy markers LC3 (green) and p62 (red). Nuclear staining with Hoechst (blue). Scalebar 50 µm. A: clear cell RCC, patient case CC05. B: Chromophobe RCC, patient case CH01. C: Papillary RCC, patient case P02. D: Oncocytoma, patient case O06. **E**: Bar plot of median LC3 intensity in tumor cells relative to normal tissue showing no significant differences between LC3 among the subtypes. Values are log_2_ transformed. **F**: Bar plot of median p62 intensity levels in tumor cells relative to normal tissue showing significant differences between all the subtypes compared to clear cell RCC group. Values are log_2_ transformed. E-F: Statistical analysis was performed with ANOVA followed by Tukey’s test for pairwise comparisons. **** = p <0,0001, *** = p <0,001, ** = p < 0,01, * = p <0,05. Non-significant (ns) comparisons are not shown.

### Tumor samples from different subtypes cluster based on protein levels of autophagy markers

Next, we asked whether the differences observed by immunostaining could also be detected at the protein level by another method. To this end, the same biobanked fresh-frozen kidney tumor samples and corresponding normal kidney tissue from each patient were analyzed by immunoblotting (Fig. S3A-G). Immunoblotting was performed with antibodies against LC3B and GABARAPL1 as markers for autophagy. Of note, these antibodies can potentially partially detect the other ATG8 family members, respectively LC3A and LC3C, and GABARAP and GABARAPL2. Taking this into consideration, we use the terms LC3 and GABARAPL1 throughout the following. The ATG8 family proteins exist as unlipidated and lipidated proteins, where lipidated ATG8 is observed as the faster migrating band on western blots, termed ATG8-II, and lipidation correlates with autophagosome levels. The slower migrating unlipidated ATG8 band is termed ATG8-I. In addition, we used antibodies to detect the autophagy cargo adaptor SQSTM1/p62, which is also a substrate for autophagy (Fig. 3A). Clear cell RCC showed significantly decreased protein levels of LC3-I, LC3-II and GABARAPL1 in tumor, aligning with immunofluorescence analysis. For chromophobe RCC and oncocytoma, levels of LC3 and GABARAPL1 were higher in tumor samples (Fig. 3B). Papillary RCC did not show a consistent pattern across these markers. SQSTM1/p62 was generally higher in tumor than normal tissue across subtypes, a striking discrepancy to the observed immunofluorescence results for clear cell RCC (Fig. S3H).

**Figure 3:**
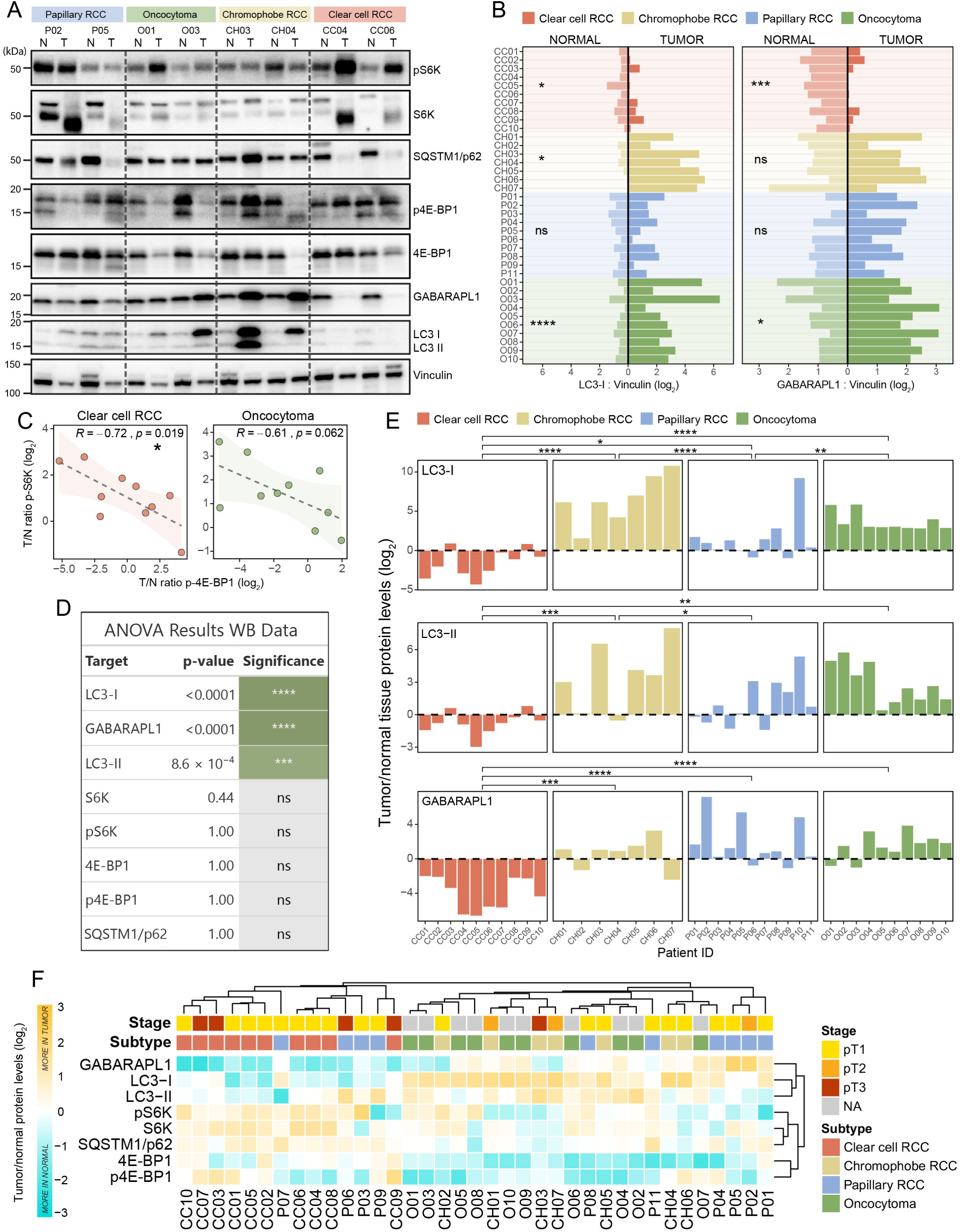
Autophagy protein levels separate clear cell RCC from other subtypes. **A**: Representative western blot for autophagy markers LC3B, GABARAPL1 and p62, and mTORC1 substrates pS6K and p4E-BP1 in kidney tumor subtypes. N = normal, T = tumor. **B**: Quantification of protein levels as determined by immunoblotting for LC3-I and GABARAPL1 (normalized to vinculin). Statistical tests are paired t-test with Benjamini-Hochberg correction for multiple comparisons. **C**: Relative tumor p-S6K versus relative tumor p-4E-BP1 levels for clear cell RCC (red) and oncocytoma (green) samples as determined by immunoblotting. Statistical test is linear regression with Pearson’s coefficient. Ribbon represents standard error of estimate. **D**: ANOVA results for log_2_ transformed tumor/normal relative values between subtypes in western blot data. **E**: Tumor/normal protein levels across subtypes for LC3-I, LC3-II and GABARAPL1. All ratio values are log_2_ scaled. Statistical comparisons are done with Tukey’s test post ANOVA. Non-significant (ns) comparisons are not shown. **F**: Heat map of tumor/normal protein levels for all targets, shown with annotations for subtype and tumor stage. All ratio values are log_2_ scaled. Higher values (orange) indicate more in tumor than normal. The heat map shows clustering of clear cell RCC samples. **** = p <0,0001, *** = p <0,001, ** = p < 0,01, * = p <0,05, ns = not significant.

mTORC1 is a major negative regulator of autophagy. As a readout of mTORC1 activity in RCC samples, we assessed phosphorylation of downstream substrates S6K and 4E-BP1. pS6K and S6K levels were generally higher in tumor across subtypes (Fig 3A, Fig. S3H). In contrast, 4E-BP showed no tumor-associated trends or clear subtype-specific clustering. Surprisingly, relative tumor levels of p-S6K negatively correlated with p-4E-BP1 in clear cell RCC, with the same trend being observed for oncocytoma (Fig. 3C), suggesting independent regulation of these mTORC1 downstream targets in these subtypes. This association was not observed in chromophobe and papillary RCC (Fig. S3I). Moreover, elevated p4E-BP levels in tumor correlated with higher tumor stage specifically in clear cell RCC (Fig. S3J), whereas no such correlation was seen for pS6K (Fig. S3K), further suggesting mTORC1 independent phosphorylation or selective regulation in clear cell RCC.

To compare differential protein levels between subtypes we calculated tumor to normal ratios for all markers across samples. ANOVA confirmed that relative levels of LC3-I, GABARAPL1 and LC3-II in tumor differed between RCC subtypes (Fig. 3D). Specifically, clear cell RCC displayed lower tumor levels of LC3-I and GABARAPL1 compared to the other subtypes (Fig. 3E). Moreover, chromophobe RCC had significantly higher LC3-II levels in tumor compared to clear cell RCC and papillary RCC (Fig. 3E). Overall, combined profiling of tumor to normal values form immunoblotting revealed strong clustering of clear cell RCC samples, while it failed to differentiate between chromophobe RCC, papillary RCC and oncocytoma (Fig. 3F).

### Differences of autophagy proteins are partially explained by transcriptional changes

Next, we wanted to investigate if observed differences in protein levels were due to differential transcriptional regulation. Therefore, we analyzed mRNA levels by quantitative PCR for the genes corresponding to the previously mentioned proteins. Additionally, we included an extended selection of genes involved in autophagy to get a broader understanding of potential transcriptional regulation of autophagy in RCC (Fig. S4A). Again, we calculated tumor expression levels relative to corresponding normal kidney for our gene panel. In general, samples failed to cluster according to subtype purely based on expression data, although two distinct groups of oncocytoma were observed (Fig. 4A). GABARAPL1 and LC3B were among the significantly differentially expressed genes between subtypes as determined by ANOVA (Fig. 4B). Our qPCR data show that levels of GABARAPL1 were higher in tumor than in normal for chromophobe RCC and oncocytoma (Fig. 4C). Moreover, relative tumor expression of LC3B was lower in clear cell RCC compared to the other subtypes (Fig. 4D). We also compared relative tumor protein level (WB) and transcriptional level (RQ) for GABARAPL1 and LC3, which confirmed that the observed differential GABARAPL1 protein levels in clear cell and chromophobe RCC could partially be explained by transcriptional changes (Fig. 4F). Interestingly, this correlation was not observed for papillary RCC where transcription of GABARAPL1 remained low compared to normal tissue, despite high protein levels in tumor samples. Similar correlation was seen for LC3; the observed higher LC3-I protein levels in chromophobe RCC and oncocytoma seem to be reflected in transcription levels of LC3B (Fig. 4G).

**Figure 4:**
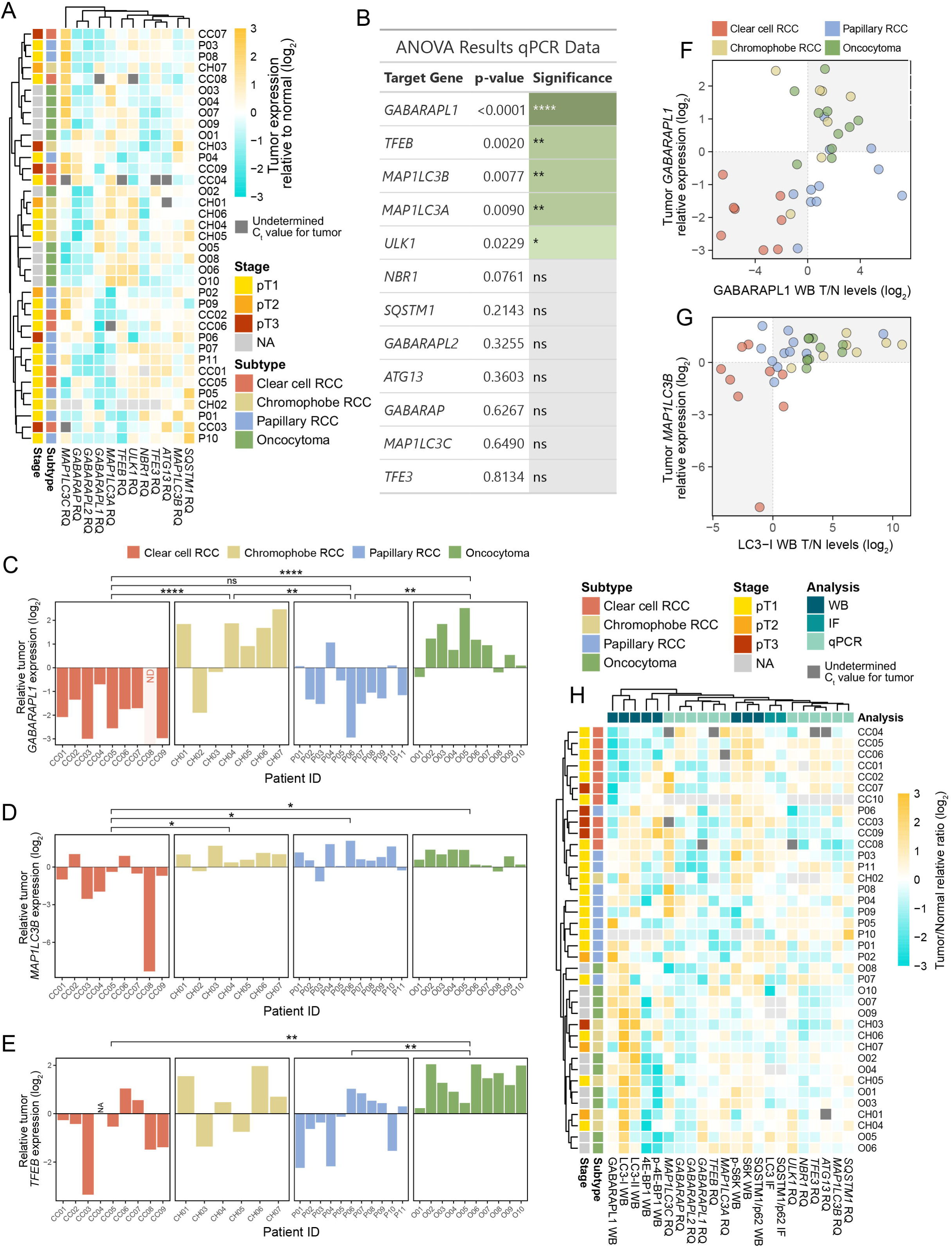
Gene transcription by itself does not recapitulate differences between kidney tumor subtypes. **A**: Heat map of mRNA levels of autophagy genes in tumor sample relative to normal tissue, shown with annotation for tumor subtype and stage. Higher values (orange) indicate more in tumor than normal. Light grey indicates missing values. Dark grey indicates Ct undetectable in tumor sample. Values are log_2_ transformed. **B**: ANOVA results for qPCR data. **C-E**: Bar plots of tumor relative expression levels for *GABARAPL1* (C), *MAP1LC3B* (D) and *TFEB* (E). Values are log_2_ transformed. ND = non-detectable expression levels in tumor sample. **F-G**: Scatter plots showing relationship between relative mRNA expression and protein amount determined by western blot for GABARAPL1 (F) and LC3B/LC3-I (G). Light gray quadrant represents area where both relative mRNA expression and protein levels are lower or higher in tumor compared to normal. **H**: Heat map with combined data from analysis of protein levels and transcriptional levels. Light gray indicates missing experimental values, whereas dark grey indicates undetermined Ct value for tumor in RT-qPCR analysis. **** = p <0,0001, *** = p <0,001, ** = p < 0,01, * = p <0,05, ns = not significant.

Next, we combined values from both protein and transcriptional level to generate a heat map of all samples (Fig. 4H). Comparing clustering of subtypes based on immunoblotting, qPCR or the combined data set by Normalized Mutual Information (NMI) scores, confirmed that transcriptional information performed poorly to differentiate tumor subtypes with an NMI score of 0.28 (Fig. S4B). However, combining expression data, immunofluorescence data, and immunoblot data improved the clustering of subtypes from 0.365 for western blot measurements only to 0.575 for the combined data set (Fig. S4B). Overall, the observed subtype-specific protein level patterns appear to be partly driven by transcriptional regulation, particularly in chromophobe RCC. Together, these findings suggest that autophagy component measurements at the protein-level capture RCC subtype specific differences where post-translational mechanisms are also likely to be involved.

### RCC cell lines show differences in autophagy flux and mTORC1 signaling

To support our findings from tissue sample experiments with functional assays, we complemented with experiments in RCC cell lines. For this we used cell lines 786-O and Caki-1 to represent clear cell RCC, A498 and Caki-2 to represent papillary RCC and HK-2, a proximal tubular cell line, as a control. Of notice, 786-O, Caki-2 and A498 harbors loss-of-function mutations in the tumor suppressor *VHL*, while Caki-1 does not. No commercially available cell lines properly represent chromophobe RCC, meaning that a cell line model for this RCC subtype is unfortunately lacking. In addition to having clear cell origin, Caki-1 cells are derived from RCC metastasis in the skin, whereas 786-O, Caki-2 and A498 originate from primary renal tumors.

To study autophagy flux and mTORC1 activity in these cell lines, they were cultured in complete medium or exposed to amino acid and serum starvation for 1-4 hours in the presence or absence of the lysosomal inhibitor BafA_1_ (Klionsky et al., 2021). In addition, we included SAR405 as an inhibitor of autophagosome formation (Klionsky et al., 2021). mTORC1 activity was assessed by phosphorylation of its downstream target S6K. 786-O, Caki-1 and Caki-2 showed the expected decrease in pS6K indicating inhibited mTORC1 activity upon starvation (Fig. S5A-C), similarly to what was observed for non-malignant HK-2 cells (Fig. S5E). As for A498, S6K remained strongly phosphorylated during starvation but decreased with BafA_1_ treatment (Fig. S5D), suggesting canonical mTORC1 hyperactivity seemingly dependent on functional V-ATPase.

LC3 lipidation and turnover of SQSTM1/p62 was assessed in the presence of nutrients and upon EBSS starvation to evaluate autophagic capacity of the different cell lines. Starvation led to an increase in lipidated LC3 in all cell lines, which was abolished by SAR405 as expected (Fig. S5A-E). Out of the four RCC cell lines, Caki-1 cells had significantly higher autophagy capacity in response to starvation, as shown by LC3 flux (Fig. 5A). SQSTM1/p62 turnover was more variable between cell lines and did not consistently reflect LC3-II lipidation levels (Fig. S5A-E). In 786-O cells, SQSTM1/p62 flux was detectable already after 1 hour of starvation, whereas this increase was not observed until 2 hours of starvation in A498 (Fig. S5F). In comparison, SQSTM1/p62 flux in HK-2 cells during starvation remained low compared to A498 throughout starvation (Fig. S5F).

**Figure 5:**
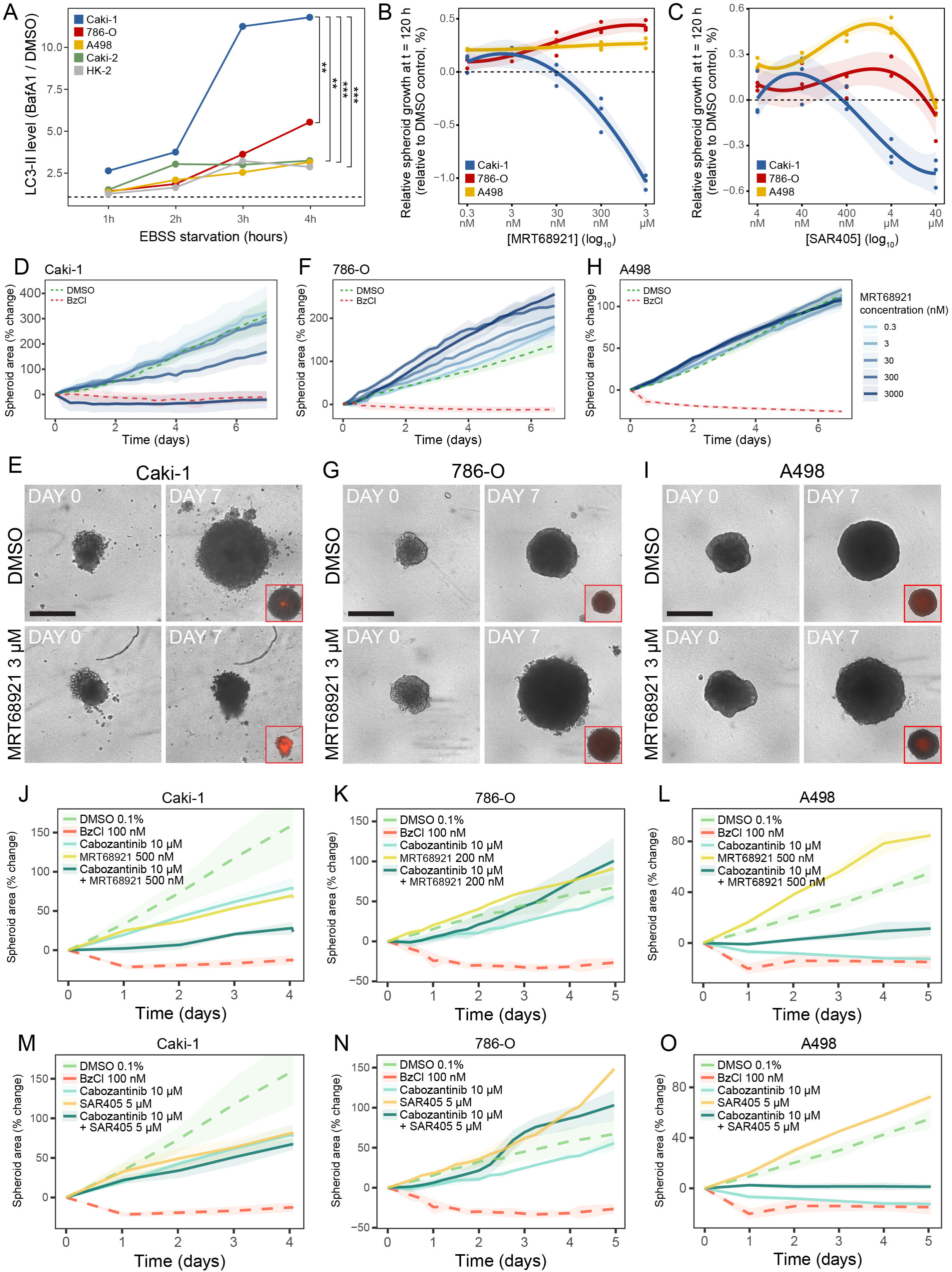
Drug treatment of RCC spheroids reveals differential response to autophagy modulation. **A:** LC3-II levels as measured by immunoblotting in RCC cell lines and normal kidney control cell line HK-2. Values are BafA_1_/DMSO ratios per hour throughout 4 hours of EBSS starvation. Horizontal dotted line represents ratio value 1. ** = p < 0,01, * = p <0,05. B, C: Spheroid growth response at t = 120 hours shown as relative growth compared to DMSO control for RCC cell lines Caki-1 (blue), 786-O (red) and A498 (yellow) after drug treatment with ULK1 inhibitor MRT68921 (B) or PI3KC3/Vps34 inhibitor SAR405 (C). Response curves are modelled with b-splines for the log_2_ ratios of the drug response values divided by DMSO, and confidence intervals represent 95% confidence. Each datapoint represents a technical replicate. Values for DMSO are obtained by fitting a Gaussian Process regression model on the growth at t=120 for all DMSO conditioned by experiment replicate and averaging them out. D, F, H: Spheroid growth shown as % change from t = 0 in response to ULK1 inhibitor MRT68921 over 7 days for cell lines Caki-1 (C), 786-O (E) and A498 (G). Negative control (DMSO vehicle) is shown as green dashed line and positive control BzCl 100 µM as red dashed line. Line represents the mean of replicate wells for a given timepoint, and shaded ribbon represents the SD across replicates. E, G, I: Brightfield images of spheroids treated with DMSO and highest concentration of ULK1 inhibitor MRT68921 (3 µM) from day 0 and day 7 of drug treatments for Caki-1 (D), 786-O (F) and A498 (H). Days are relative to when drugs were added to spheroid culture. Inserts in red frame: Cytotox Red signal indicating cell death in spheroids at day 7 of drug treatment. Scale bar 500 µm. J-L: Spheroid growth shown as % change from t = 0 in response to combination treatment of 10 µM cabozantinib and ULK1i MRT68921. Note that ULK1i concentration for 786-O (L) is 200 nM, while it is 500 nM for Caki-1 (K) and A498 (M). M-O: Spheroid growth shown as % change from t = 0 in response to combination treatment of 10 µM cabozantinib and 5 µM SAR405 for Caki-1 (N), 786-O (O) and A498 (P). Negative control (DMSO) is shown as green dashed line and positive control BzCl 100 µM as red dashed line. Line represents the mean of replicate wells for a given timepoint, and shaded ribbon represents the SD across replicates.

### Different RCC cell lines exhibit opposing response to autophagy modulation

We next investigated whether different RCC cell lines had differential response to autophagy modulation by drugs, as they showed varied autophagy response to starvation and under basal conditions. To do this, we established 3D culture spheroids to better mimic cellular environment and recapitulate the solid tumor (Jensen & Teng, 2020). Spheroids were successfully established in 2.5% Basement Membrane Extract (BME) gel for three out of the four RCC cell lines: 786-O, A498 and Caki-1 (Fig. S5G). Despite improved spheroid formation after addition of BME, Caki-2 and HK-2 generally displayed inconsistent spheroid formation and limited growth and was excluded from drug assays dependent on spheroid growth as readout (Fig. S5H). RCC spheroids were treated with drugs known to modulate autophagy in various ways. Torin 1 was used to increase autophagy levels by blocking MTOR kinase activity. All cell lines were sensitive to Torin 1 treatment (Fig. S5I). MRT68921 and SAR405 were used to inhibit autophagy induction as they are inhibitors of ULK1 and Vps34, respectively. Spheroid size was measured over time, and Cytotox Red was added to assess viability enabling differentiation between growth restriction and cell death. Strikingly, we observed different responses to autophagy inhibition across the three RCC spheroid models (Fig. 5B-C). Caki-1 spheroids were sensitive to ULK1 inhibition and displayed both reduced growth and increased cell death (Fig. 5D-E). In contrast, 786-O and A498 spheroids were not sensitive to autophagy inhibition by ULK1i (Fig. 5F-I). In fact, 786-O spheroids displayed increased spheroid growth upon ULK1 inhibition in a concentration dependent manner (Fig. 5F-G). Similar concentration dependent growth increase was also observed with SAR405 treatment up to 4 µM for 786-O and A498 spheroids, but these cell lines displayed a biphasic response with cytotoxic effects at 40 µM SAR405 (Fig. 5C).

Next, we asked if autophagy inhibition could affect the outcome of treatment efficacy. Cabozantinib is a tyrosine kinase inhibitor used in first-line therapy for advanced RCC. In Caki-1 spheroids, combining 10 µM cabozantinib with 500 nM MRT68921 reduced spheroid growth more than individual treatments alone (Fig. 5J). 786-O spheroids were less sensitive to cabozantinib treatment, and the addition of MRT68921 had no effect on the spheroid growth (Fig. 5K). Strikingly, the addition of ULK1 inhibitor to the cabozantinib sensitive cell line A498 worsened the outcome in regards of spheroid growth (Fig. 5L). The lack of potentiating cabozantinib cytotoxic effects on spheroid growth upon autophagy inhibition was confirmed with SAR405, which displayed similar response patterns for 786-O and A498 as for ULK1 inhibition (Fig. 5N-O), although the combinatorial effect for Caki-1 was not observed with Vps34 inhibition (Fig. 5M). To gain insight into autophagy-related differences between the RCC cell lines, we explored their expression levels of selected autophagy-related genes in the public CCLE database. Importantly, Caki-1 did not show elevated expression of ULK1, ULK2 and PIK3C3 compared to 786-O, indicating that transcriptional upregulation of MRT68291 and SAR405 drug targets was unlikely to account for the increased sensitivity (Fig. S5J). These results suggest that sensitivity to autophagy modulation in RCC is not limited to histological subtype, but that molecular profiles and tumor stage might impact the response.

## Discussion

Autophagy has been shown to contribute to cancer development, including tumor progression and treatment response (Guilbaud et al., 2026). Understanding the role of autophagy in specific cancer types may have benefits for therapeutical approaches. In this study, we aimed to get a better understanding of autophagy in renal cell carcinoma. We have demonstrated that protein levels of autophagy markers can be used to distinguish clear cell RCC from non-clear cell RCC subtypes. Moreover, combining protein levels with transcriptional data distinguishes clear cell RCC and papillary RCC from chromophobe RCC and oncocytoma. We therefore suggest that protein levels of autophagy markers can help to discriminate between kidney tumor subtypes. Similar conclusion were drawn in a study discriminating renal cell carcinoma subtypes based on immunohistochemistry images of key autophagy proteins (He et al., 2020). Aligning with our data, they found that downregulation of LC3B characterized clear cell RCC. In contrast to the findings of us and others, a few studies have demonstrated an increase in LC3-I and LC3-II in clear cell RCC tumors compared to normal tissue as determined by immunoblotting (Guo et al., 2019; Patergnani et al., 2025). Moreover, He *et al* showed that staining against ULK1, ATG5, ATG16L1, LC3B and p62 allowed for differentiation between clear cell RCC, chromophobe RCC and normal tissue samples (He et al., 2020). In our study, we validate this discrimination in paired tumor-normal samples from fresh-frozen tissue and complement with subtype specific patterns for papillary RCC and benign oncocytoma.

Besides classification, we sought insight into how autophagy flux is affected in RCC. This is a difficult question to answer solely based on non-viable tissue samples. Immunofluorescence revealed low levels of LC3 and p62 in clear cell RCC tumor samples, suggesting high autophagy turnover of these substrates. However, the findings for p62 in clear cell RCC were not recapitulated by our immunoblotting analysis, therefore conclusions regarding high flux in clear cell RCC samples relative to the other subtypes should be approached with caution. Since we mainly identify changes in proteins of the ATG8 family (LC3 and GABARAPL1), which are all turned over by the autophagy pathway, these differences may indicate different autophagic activity between RCC subtypes. To our knowledge, no studies have specifically assessed autophagy flux across RCC subtypes, although existing studies give some indications. One line of indirect line of evidence comes from a study of kinase activities in RCC, which identified higher activity of class I PI3K in chromophobe, papillary and sarcomatoid renal cancer. Oppositely, in clear cell RCC, the PI3K pathway appeared strongly inhibited (Tahiri et al., 2022). Considering that PI3K is a positive regulator of mTORC1, these results could potentially point to a downregulation of autophagy in chromophobe, papillary and sarcomatoid renal cancer and an upregulation of autophagy in clear cell RCC. We used samples from the same biobank as Tahiri *et al*., although not exactly the same subset, and found that autophagy markers LC3B and GABARAPL1 are decreased in clear cell RCC both on mRNA and protein levels.

However, since autophagy is a dynamic process, single measurements of protein levels cannot reliably indicate whether autophagy is up- or down-regulated. Lower levels of proteins that are turned over by autophagy could result from higher autophagic turnover or simply lower gene expression. To try to address this across RCC subtypes, we assessed autophagy flux in RCC cell lines representing some of the subtypes. Functional assays revealed that the metastatic clear cell RCC cell line Caki-1 had higher autophagic capacity during starvation compared to HK-2 control cells, accompanied by higher basal LC3 flux. Aligning with this, induction of epithelial to mesenchymal differentiation in RCC cell line A498 has been shown to upregulate autophagy flux (Singla & Bhattacharyya, 2017). Notably, since no representative cell line model for chromophobe RCC exists, we were unable to evaluate flux for this subtype.

Elevated levels of p-S6K were observed in clear cell RCC, which potentially indicate dual activation of autophagy and mTORC1 in clear cell RCC. mTORC1 hyperactivity is established in VHL-deficient clear cell RCC and correlates with poor outcome (Robb et al., 2007). Consistent with this, we found that p-4E-BP1 tumor levels were higher in stage 3 clear cell RCC tumors. This pattern was not observed for p-S6K, suggesting mTORC1-independent regulation of 4E-BP1 phosphorylation or selective regulation of this substrate by mTORC1. Interestingly, we observed that mTORC1 was not fully inactivated upon starvation in A498 cells, suggesting decoupling of nutrient sensing to canonical mTORC1 activity in papillary RCC which warrants further research.

The observed differences between subtypes are not completely determined by transcriptional changes, as illustrated by the discordance between mRNA and protein levels in papillary RCC. This suggests that some of the observed differences are due to post-translational regulation and likely differences in flux through the autophagy pathway itself. Similarly, He et al. (2020) did not find direct correspondence between mRNA levels of autophagy genes and their corresponding protein levels (He et al., 2020). Another study identified higher transcriptional levels of autophagy markers *GABARAP* and *SQSTM1* in primary clear cell RCC tumors compared to normal kidney tissue (Radovanovic et al., 2020). They also found that higher levels of these markers were associated with earlier tumor stage, smaller tumor size and better survival, suggesting that transcriptional induction of autophagy in clear cell RCC might slow tumor progression and metastasis.

We consistently observed low expression of *GABARAPL1* in clear cell and papillary RCC, and high expression of *GABARAPL1* in chromophobe RCC. Lower *GABARAPL1* expression was associated with poorer overall survival for clear cell RCC in TCGA database. Given that clear cell and papillary RCC generally are more aggressive subtypes, we can speculate that *GABARAPL1* expression can serve as risk factor across RCC subtypes. Future studies will have to elucidate the potential prognostic role.

Our RCC spheroid experiments showed increased cell death and reduced cell growth for Caki-1 upon ULK1 inhibition alone and in combination with cabozantinib, suggesting that autophagy might be involved in tumor growth. Interestingly, 786-O spheroids increased in size upon ULK1 inhibition, which could indicate that autophagy might act as a tumor suppressor in this context. Several cell line characteristics could explain this discrepancy in response. Caki-1 represents metastatic clear cell RCC, whereas 786-O is considered as primary clear cell RCC (Brodaczewska et al., 2016). Moreover, while 786-O and A498 carry loss-of-function *VHL* mutations, Caki-1 retains functional VHL. This indicates that molecular profiles could inform whether autophagy inhibition or activation would be the potential therapeutical approach in RCC. Future studies are needed to determine whether VHL status is essential for autophagy dependency. Such molecular dependencies have been shown by others, for example that *BAP1* mutant RCC specifically is sensitive for autophagy suppression (Vega-Rubin-de-Celis et al., 2025). Moreover, we show that combining autophagy inhibition with tyrosine kinase inhibitor potentiated the growth of 786-O and A498 spheroids, in stark contrast to reports in the same cell lines or other clear cell RCC cell lines from primary tumors (Zheng et al., 2015; Patergnani et al., 2025). Importantly, whereas these previous studies assessed viability under conventional 2D culture conditions, our model evaluated 3D spheroid growth, which may more closely resemble tumor growth *in vivo*.

Autophagy protein levels could serve beyond differentiation between subtypes as prognostic biomarkers or aid in supporting treatment decisions. Interestingly, it was previously found that ULK1 expression levels correlated with progression-free survival of patients with metastatic RCC (Nishikawa, Miyake, Bing, et al., 2015). The same study suggested that autophagy markers could be considered as a parameter in selecting patients who will benefit from mTOR inhibitors. The same group showed that expression of Beclin 1 was predictive of reoccurrence in patients with clear cell RCC undergoing radical nephrectomy in a retrospective study (Nishikawa, Miyake, Liu, et al., 2015). Moreover, high expression of REDD1, a positive regulator of autophagy, predicts response to autophagy inhibition in RCC (Carew et al., 2025). Potential prognostic value of autophagy related proteins should be further studied at larger scale in prospective clinical studies.

How autophagy is affected in different cancer types can potentially help to find better treatment options for patients that suffer from therapy resistance. We have demonstrated that autophagy proteins differ between subtypes of RCC, which may reflect differences in autophagic activity. We have also demonstrated that spheroids derived from RCC cell lines display different responses to autophagy inhibition, indicating that the therapeutic potential is context dependent. This provides a baseline to explore how autophagy is involved in the development of RCC and whether it can be a potentially target in novel RCC therapies.

### Limitations of this study

The main limitation of this study is that our analysis methods were limited to bulk measurements. Although obvious necrotic areas were not included, our analysis does not differentiate between contributions from epithelial and stromal cells. Moreover, we have relatively few samples in our cohort. Larger sample sets might improve differentiation between subtypes, especially for evaluating trends across tumor stages. Our small samples size disabled us from performing supervised machine learning classifications like described by others (He et al., 2020).

An inherited limitation of utilizing patient tissue is the inability to perform genetic or pharmacological perturbations to validate detection of targets with antibody-based methods. Consequently, evaluation of protein target levels in patient tissue samples is more challenging and the findings warrant careful interpretation. In this study, this is exemplified by the discrepancies between SQSTM1/p62 levels observed in clear cell RCC on immunofluorescence and immunoblotting.

We aimed to complement biobank patient samples with functional flux assays and drug treatments in RCC cell lines spanning across subtypes. The main finding in patient samples was that clear cell and papillary RCC differ from chromophobe RCC in autophagy protein profiles. However, the lack of a well-characterized, commercially available chromophobe cell line prevents functional validation for this subtype. This limits the interpretations of our findings regarding autophagy dependence across all three RCC subtypes.

We used single, sub-inhibitory concentrations of autophagy modulating drugs to assess potential combinatorial effects with tyrosine kinase inhibitor cabozantinib. We appreciate that complete concentration ranges of both drugs are necessary to determine true synergistic effects. Furthermore, drug assays could be supplemented with genetic perturbation of autophagy, for example by ATG7 or ULK1 knock-down or knock-out, to strengthen the evidence that the observed effects are due to autophagy inhibition and not other pathways affected by the drugs.

## Materials and methods

### Patient samples

We used samples from the Akershus University Hospital kidney cancer biobank, which contains consecutively collected nephrectomy samples from 2013 to 2019. The biobank contains fresh-frozen samples of normal tissue and up to three tumor tissue samples from approximately 171 clear cell, 44 papillary, 13 chromophobe, 33 oncocytoma and 30 mixed/other RCC cases. From these we randomly selected 10 samples of clear cell, 7 of chromophobe and 11 of papillary RCC as well as 10 oncocytoma samples that were used in this study. An experienced pathologist assessed hematoxylin/eosin staining of tumor and normal tissue collected from each patient to ensure that these were relatively homogenous (containing either tumor or normal tissue), with minimal necrosis and free from any signs of damage. Samples were inspected morphologically to confirm the reported histological classification

### Ethical approval

The study was conducted in accordance with the principles of the Declaration of Helsinki. The biobanked samples were collected upon informed and written consent (REK 2012/771). The use of samples in the biobank has been approved for this project by the Regional Committee for Medical Research Ethics (REK South-East Norway, #48926).

### Tissue lysis

Patient samples were cryosectioned at –20 °C. The cryostat was disinfected with 1h UV-light between each session of cryosectioning and cleaned with absolute ethanol between each patient to avoid contamination. Roughly 15 x 10 µm sections were added to pre-cooled Eppendorf tubes with RIPA-buffer (10mM Tris pH 8, 140 mM NaCl, 1 mM EDTA, 0,1% Na deoxycolate, 0,1% SDS, 1% Triton X-100) supplemented with protease (Merck Life Science #539131-1VL) and phosphatase (Roche #04906837001) inhibitors. These lysates were centrifuged in QIAshredder columns (Qiagen #79656) at 1200 rpm followed by sonification and centrifugation for 15 min at 12000 rpm in 4° C. Protein concentrations were measured with BCA assay (Thermo Scientific Pierce BCA Protein Assay Kit #23227).

### Western blot

Equal amounts of protein samples were mixed with 4x Nu-Page-buffer (Invitrogen #NP0008), 1M DTT (Sigma-Aldrich, #D0632) and water. Samples were boiled for 10 min at 95 °C, followed by loading on 4-20% SDS-polyacrylamide Criterion™ TGX™ Precast Gel (BioRad #567-1095). Proteins were separated by gel electrophoresis at 80V-150V. The LF-PDVF membrane (BioRad Trans-Blot Turbo RTA Transfer Kit #1704275) was activated with absolute EtOH both before semi-dry transferring (BioRad Trans-Blot Turbo^TM^ Tansfer system) and prior to blocking by air drying. Then the membranes were incubated with primary antibodies in BSA (Sigma-Aldrich, #A2153) or milk (Sigma-Aldrich #70166-500G) at 4 °C overnight. The next day they were washed with 1xTBS-Tween (0,1%) (Sigma-Aldrich, #P1379) 4×5 min before incubation with HRP-conjugated secondary antibodies in 5% milk. Then they were washed 4×5 min and developed with Dura (Thermo Scientific Supersignal^TM^ West Dura Extended Duration Substrate #34076). The images were developed with Sapphire™ Biomolecular Imager (Azure biosystems). Quantification was performed using ImageLab 6.1 Software. The intensity of each band was measured, and the background intensity was subtracted using the Global Subtraction Method. The volume value for each band was normalized to the level of a housekeeping protein (vinculin or actin). Patient sample P10 was omitted from western blot analysis due to inadequate protein lysate quality.

### Cell culture and cell lines

Several RCC cell lines were used as representative of the subtypes. Cells were cultured in the following media: for clear cell RCC we used 786-O cells (ATCC, CRL-1932) cultured in RPMI (Sigma #R2405), and Caki-1 (ATCC, HTB-46) and Caki-2 (ATCC, HTB-47) cells cultured in McCoy’s 5a Medium Modified (Biowest #L0210-500). A498 (ATCC, HTB-44) cells representing papillary subtype were cultured in Eagle’s Minimum Essential Medium (ATCC #30-2003). HK2 (ATCC, CRL-2190, Lot# 70041172) cells derived from kidney were used as control and cultured in Dulbecco’s Modified Eagle Medium/F12 (Gibco #31331). All media were supplemented with 10% inactivated FBS (Sigma-Aldrich #F7524) and 1% Penicillin/Streptomycin (Gibco #15140122). All cells were maintained at 37 °C and 5% CO_2_. Cells were routinely checked for mycoplasma contamination and found to be mycoplasma free.

### Autophagy flux assessment in RCC cell lines

LC3 and p62 flux as well as mTORC1 signaling in RCC cell lines were analyzed with starvation in combination with drug treatment. Cells were starved in EBSS (Thermo Fisher Scientific, #24010043) for 1-4 hours and treated with either vehicle (0.1% DMSO, Sigma #D8418), 100 nM BafA1 (Enzo life sciences #BML-CM-110-0160) or 5 µM SAR405 (Selleckchem #S7682). Cells were lysed with RIPA buffer and collected by scraping in microcentrifuge tubes with lysis buffer. Western blot was performed according to the protocol above except for the step with Qiashredder.

### Antibodies used for immunoblotting

LC3B (Cell Signaling Technologies #2775) 1:1000 in 5% BSA, p-S6K Thr389 (Cell Signaling Technologies #9205) 1:1000 in 5% BSA, S6K (Cell Signaling Technologies #9202) 1:1000 in 5% BSA, p62 (Abnova #H00008878-M01) 1:1000 in 5% milk, GABARAPL1 (Proteintech #18721-1-AP) 1:1000 in 5% BSA, 4E-BP 1:1000 (Cell Signaling Technologies #9644), p-4E-BP1 Thr37/46 (Cell Signaling Technologies #2855) 1:1000 in 5% BSA, Vinculin (Sigma Aldrich #V9131) 1:10 000 in 5% BSA, beta-actin (Abcam #ab8227) 1:1000 in 5% BSA. HRP-goat-anti mouse (Jackson #115-035-003) and HRP-goat-anti rabbit (Jackson #111-035-144).

### RNA extractions

Patient samples were cryosectioned as described above and roughly 15 x 10 µm sections were added to pre-cooled Eppendorf tubes with lysis buffer (Buffer RLT Plus). RNA was extracted with the All-prep DNA/RNA kit (Qiagen #80284) according to manufacturer’s description. Nucleic acid concentrations were measured with NanoDrop 2000 (Thermo Fisher).

### RT-qPCR

For qPCR analysis, reverse transcription was performed on up to 1000 ng RNA using the iScript^TM^ cDNA Synthesis Kit (BioRad #1708890). Two technical replicates per sample were used for each experiment. qPCR reactions were performed with SYBRgreen (Thermo Fisher #4385617) with final primer concentration of 0.2 µM. The total amount of cDNA was 1.25 ng per reaction. The quantification of target genes was determined using the Ct method with multiple reference genes to stabilize gene expression data (Riedel et al., 2014). Briefly, the Ct (threshold cycle) values of target genes were normalized to two reference genes (TBP and Cyclophilin A (chromophobe), TBP and 18srRNA (clear cell, papillary and oncocytoma) or TBP and GAPDH (patient IDs P05, O01, CH02)) where ΔCt = Ct_target_ – Ct_ref_, and then compared with a calibrator sample (normal tissue) where ΔΔCt = ΔCt_Sample_ -ΔCt_Calibrator_. Relative expression (RQ) was calculated with the formula RQ = 2^-ΔΔCT^.

### Primers used in this study

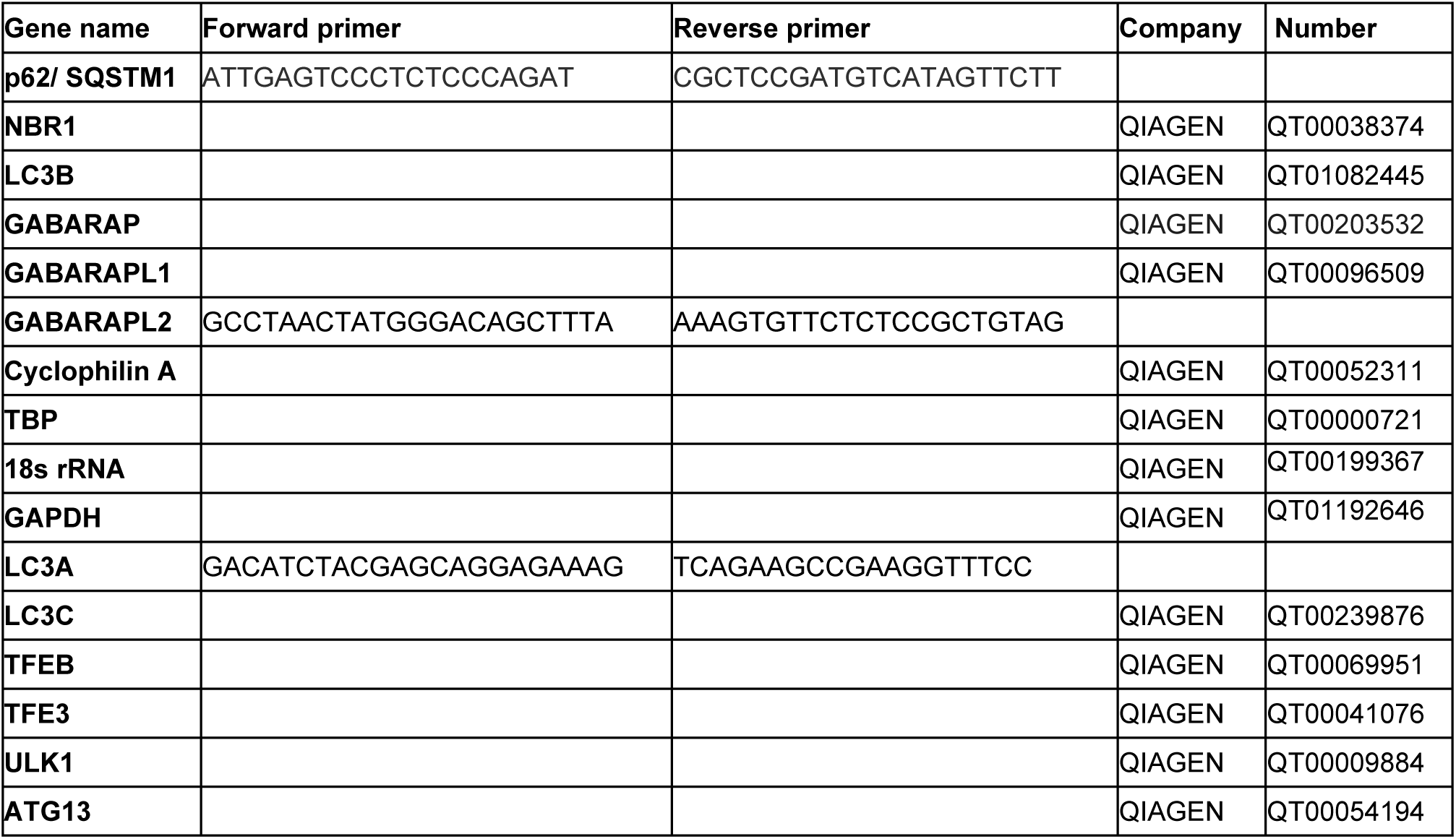

### Immunofluorescence on tissue sections

Patient samples were cryosectioned at 5 µm and collected on glass slides on dry ice before fixation with 4% PFA (Polysciences, #18814-20) for 10 minutes, followed by a rinse with PBS. The slides were then blocked in 5% BSA in PBS-T (0,1% Tween-20) for 30 minutes. Then the slides were stained with primary antibodies anti-LC3B (MBL #PM036 1:1000), anti-p62 (MBL #610833 1:300) in PBS-T with 5% BSA at 4 °C overnight. The next day the slides were washed 3x 5 min with PBS-T, followed by secondary staining with Alexa Fluor 647-donkey-anti-rabbit (Jackson #711-605 -152) and Alexa Fluor 568-donkey-anti-mouse (Thermo Fisher #A10037) in PBS-T with 5% BSA for 3h at room temperature. For nuclear staining, the slides were incubated with 1 µg/ml Hoechst 33342 (Thermo Scientific, #62249) in PBS-T with 5% BSA for 10 min. Afterwards they were rinsed with PBS-T and mounted with Prolong^TM^ Diamond Antifade Mountant (Invitrogen #P36961). Slides were imaged on VS200 Slide Scanner (OLYMPUS) equipped with a X-Cite XYLIS LED light source (Excelitas tech) and a X-Line Apo 40x NA0.95 Air objective. Images were acquired with a Z stack of 7 µm. Images were analyzed with QuPath (v0.5.0) (Bankhead et al., 2017).

### Data analysis for patient material

Data analysis was conducted in R (v 3.4.2). Heatmaps were made with the *pheatmap* package (v 1.0.12) using complete linkage hierarchical clustering with the Euclidean distance measure and row scaling of log_2_ transformed values. Normalized Mutual Information (NMI) was calculated to assess patient sample heatmap clustering algorithm for k=4 using the *aricode* package (v 1.0.3). All data was assumed normally distributed after log_2_ transformation. Multiple comparison corrections were performed when pairwise comparing more than two samples with details stated in figure legends.

### Spheroid drug treatment

2000 cells were seeded in 100 µl complete media (as described above) containing 3% Matrigel® Matrix Basement Membrane (GF Reduced) - Phenol-Red Free (# 356231 Lot# 312800) or Cultrex™ Reduced Growth Factor BME Type 2 (# 3533-005-02 Lot# 1699846) in Corning® 96-well Clear Round Bottom Ultra-Low Attachment microplate (#7007) or Sarstedt BIOFLOAT™ 96-well microplate (#83.3925.400). The plate was centrifuged at 100 x g for 10 minutes to avoid formation of multiple spheroids before incubation at 37℃ to allow for spheroid formation. After 3 days, 100 µl of complete media containing drugs at 2x of final concentration were added to each well. Drugs used in this study were Torin1 (Tocris, #4247), MRT68921 (Sigma Aldrich, #SML1644), SAR405 (Selleckchem, #S7682) and Cabozantinib (Selleckchem, #S1119). DMSO concentrations were matched for all drug dilutions to a final concentration of maximum 0.4 %. Negative control wells were treated with DMSO vehicle. Positive control for cell death was 100 µM benzethonium chloride (BzCl) (Santa Cruz Biotechnology, #sc-239299). IncuCyte® Cytotox Red Fluorescent Dye (Sartorius Essenbio, #4632, Lot#2305004113) was added in a final concentration of 200 nM to monitor cell death. Spheroid growth was monitored every 6^th^ hour for 5-7 days with IncuCyte® S3 Live Cell Analysis Instrument (Sartorius Essenbio). Spheroids were imaged with 4x objective. The spheroid area and Cytotox Red intensity was measured using the IncuCyte® Spheroid module.

### Data analysis for spheroid drug treatments

Data analysis was conducted in R (v 4.6.1). For drug concentration response curves in Fig 5A-B, outlier removal was performed utilizing a smoothing moving average. To estimate normalized drug treatment effects relative to DMSO controls we first fitted a longitudinal model over the different DMSO curves with additive Gaussian Process regression using the *lgpr* package (v. 1.2.5) (Timonen et al., 2021). Log_2_ ratios were then computed of the observed drug response values vs the estimated DMSO curves. Curves and 95% confidence intervals for each cell line were generated using the b-splines implementation from the loess function of the base R package.

### CCLE expression data for RCC cell lines

RNA-seq gene expression data was extracted as transcriptions per million (TPM) values from the CCLE database (Barretina et al., 2012). Expression was reported relative to HEKTE expression, the only normal kidney cell line present in CCLE dataset. Heatmap for gene expression data was made with the *ComplexHeatmap* package (v 2.28.0) in R (v 4.6.1) (Gu et al., 2016). Clustering for the heatmap is done with hierarchical clustering with Euclidean distances.

## Acknowledgements

We thank Martina Landschoof Skrede for kindly providing training for cryosectioning. We thank Oslo University Hospital core facility for Advanced Light Microscopy Montebello node and Molmed Imaging Platform. F.K. and H.U. are part of The Medical Student Research Program (MSR) at the Faculty of Medicine, University of Oslo, and were supported by grant 271555/F20 from the Research Council of Norway and running costs from the Faculty of Medicine. H.K. was supported by grant 300748 from the Research Council of Norway, grants 2017062 and 2022019 from the South-Eastern Norway Regional Health Authority and by the European Union (ERC, FINALphagy, 101039174). FINALphagy is a project in the Centre for Digital Life Norway, which is supported by the Research Council of Norway’s grant 320911. J.M.E. was supported by South-Eastern Norway Regional Health Authority (HSØ grants 2017064, 2018012, 2019049 and 2019096), the Norwegian Cancer Society (grants 182524 and 208012) and the Research Council of Norway through research grants 261936, 294916 and 314811, and from the European Union’s Horizon 2020 research and innovation programme under grant agreement No. 847912 (RESCUER). This work was partly supported by the Research Council of Norway through its Centre of Excellence funding scheme, project number 262652.

## Author contributions

Feyza Kasikci: Formal analysis, Investigation, Writing—original draft, Writing—review & editing, Visualization.

Julie Aarmo Johannessen: Formal analysis, Investigation, Writing—original draft, Writing— review & editing, Visualization.

Eva Bassols-Citores: Formal analysis, Writing—original draft, Writing—review & editing, Visualization

Maria Frison: Investigation, Writing—review & editing

Siri Andresen: Investigation, Writing—review & editing

Elise Komttebed: Investigation, Writing—review & editing

Frode S. Nilsen: Resources, Writing—review & editing,

Stig Müller: Resources, Writing—review & editing,

Karol Axcrona: Resources, Writing—review & editing,

Leiv Rønneberg: Writing—review & editing, Supervision

Manuela Zucknick: Writing—review & editing, Supervision

Lorant Farkas: Formal analysis, Investigation, Resources, Writing—review & editing, Visualization.

Jorrit M. Enserink: Writing—review & editing, Supervision.

Helene Knævelsrud: Conceptualization, Formal analysis, Investigation, Writing—original draft, Writing—review & editing, Supervision, Project administration, Funding acquisition.

## Declaration of interests

The authors declare that they have no competing interests.

**Table 1:** Overview of patients included in this study. Histology of RCC, age and sex of patients and stage and grade of tumors are included in the table. Chromophobe = CH, Clear cell = CC, Papillary = P, Oncocytoma = O.

| **Patient ID** | **Histology** | **Age** | **Sex** | **Stage** | **Grade** |
|---|---|---|---|---|---|
| CC01 | Clear cell | 72 | F | pT1b | 3 |
| CC02 | Clear cell | 69 | M | pT1b | 2 |
| CC03 | Clear cell | 78 | M | pT3a | 3 |
| CC04 | Clear cell | 40 | M | pT1b | 2 |
| CC05 | Clear cell | 40 | F | pT1a | 2 |
| CC06 | Clear cell | 59 | F | pT1a | 2 |
| CC07 | Clear cell | 68 | F | pT3a |  |
| CC08 | Clear cell | 81 | F | pT1a (recidive) |  |
| CC09 | Clear cell | 78 | M | pT3a |  |
| CC10 | Clear cell | 67 | M | pT1a |  |
| CH01 | Chromophobe | 55 | M | pT2b | Not relevant |
| CH02 | Chromophobe | 43 | F | pT1a |  |
| CH03 | Chromophobe | 71 | M | pT3aN2 |  |
| CH04 | Chromophobe | 52 | M | pT1a |  |
| CH05 | Chromophobe | 61 | F | pT1b |  |
| CH06 | Chromophobe | 47 | F | pT1a | Not relevant |
| CH07 | Chromophobe | 70 | F | pT2b | Not relevant |
| O01 | Oncocytoma | 74 | F | Not relevant | Not relevant |
| O02 | Oncocytoma | 80 | M | Not relevant | Not relevant |
| O03 | Oncocytoma | 71 | F | Not relevant | Not relevant |
| O04 | Oncocytoma | 62 | F | Not relevant | Not relevant |
| O05 | Oncocytoma | 72 | M | Not relevant | Not relevant |
| O06 | Oncocytoma | 70 | M | Not relevant | Not relevant |
| O07 | Oncocytoma | 81 | F | Not relevant | Not relevant |
| O08 | Oncocytoma | 62 | F | Not relevant | Not relevant |
| O09 | Oncocytoma | 67 | M | Not relevant | Not relevant |
| O10 | Oncocytoma | 56 | M | Not relevant | Not relevant |
| P01 | Papillary | 61 | M | pT1a | 3 |
| P02 | Papillary | 57 | M | pT2a | 3 |
| P03 | Papillary | 49 | M | pT1a | 2 |
| P04 | Papillary | 66 | F | pT1a | 3 |
| P05 | Papillary | 60 | M | pT1a | 3 |
| P06 | Papillary | 68 | M | pT3a | 3 |
| P07 | Papillary | 60 | M | pT1a | 2 |
| P08 | Papillary | 64 | M | pT1a | 2 |
| P09 | Papillary | 67 | M | pT1a | 2 |
| P10 | Papillary | 81 | F | pT1bN1 | 3 |
| P11 | Papillary | 59 | M | pT1a | 3 |

## Abbreviations

BafA_1_: Bafilomycin A_1_
BCA: bicinchoninic acid
BSA: bovine serum albumin
DMSO: dimethyl sulfoxide
DTT: dithiothreitol
EDTA: etylendiamintetraacetat
FBS: fetal bovine serum
HRP: horseradish peroxidase
LF-PDVF: low-fluorescence polyvinylidene fluoride
MAP1LC3B: Microtubule-associated proteins 1A/B light chain 3B
mTORC1: mammalian target of rapamycin complex 1
PenStrep: penicillin-streptomycin
RCC: renal cell carcinoma
RIPA buffer: radioimmunoprecipitation assay buffer
SDS: sodium dodecyl sulfate
SQSTM1: Sequestosome 1
Tris: tris(hydroxymethyl)aminomethane
ULK1: Uncoordinated 51-like kinase

