## Supplemental figures for "Subtype-specific autophagy profiles in renal cell carcinoma"

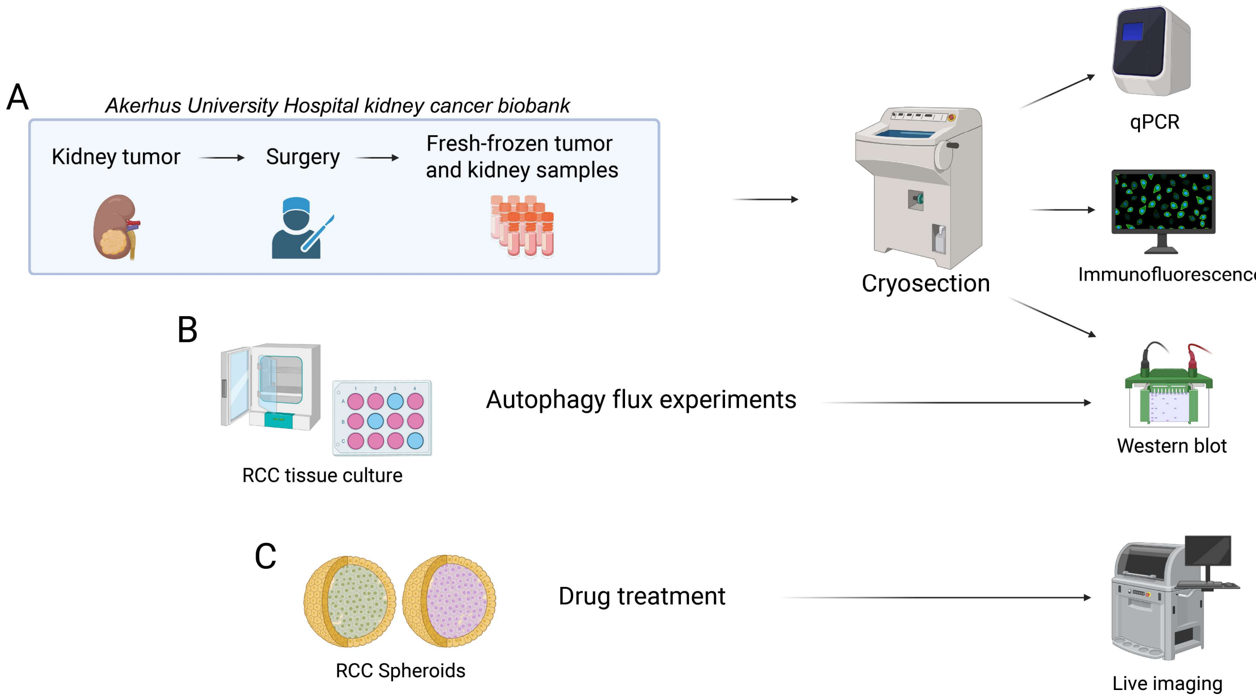


**Supplementary figure 1: Illustration of methods used in this project**. **A**: Kidney tumor and normal biopsies were taken from surgically removed kidney tumors and biobanked as fresh-frozen tissue. A subset of these samples was used in this study. **B**: Autophagy flux experiments were performed on cell lines Caki-1, Caki-2, 786-O, A498 and HK-2. **C**: RCC cell line spheroid cultures were exposed to autophagy-modulating drug treatments, both as single treatment and combination treatment with the tyrosine kinase inhibitor cabozantinib. Illustration was made with BioRender.

A

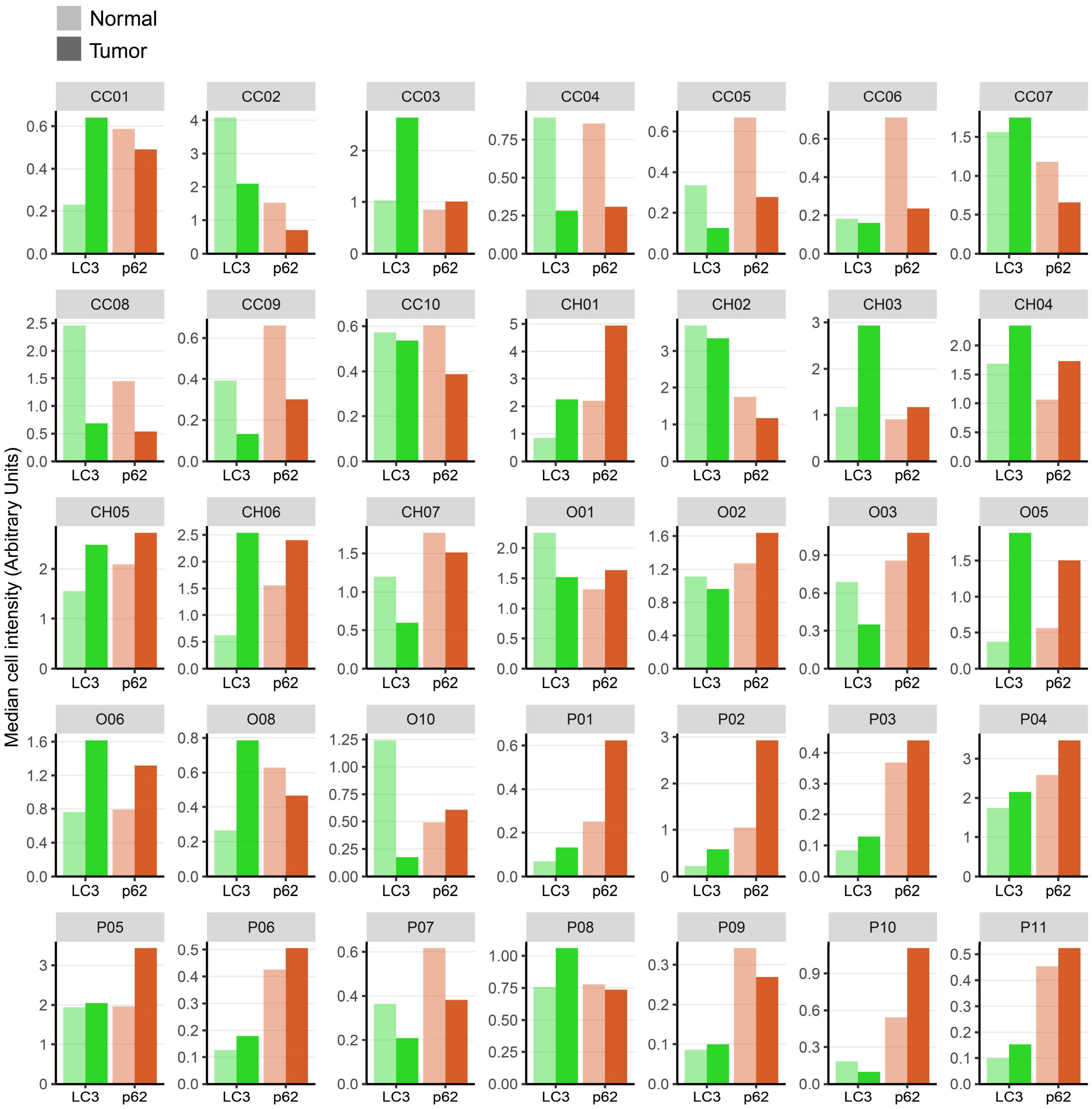


**Supplementary figure 2: Raw values for all samples included in immunofluorescence for this study.** **A**: Median cell intensity of LC3 (green) and SQSTM1/p62 (red) in tumor versus normal tissue across all samples. Normal samples are shown with more transparency than tumor sample.

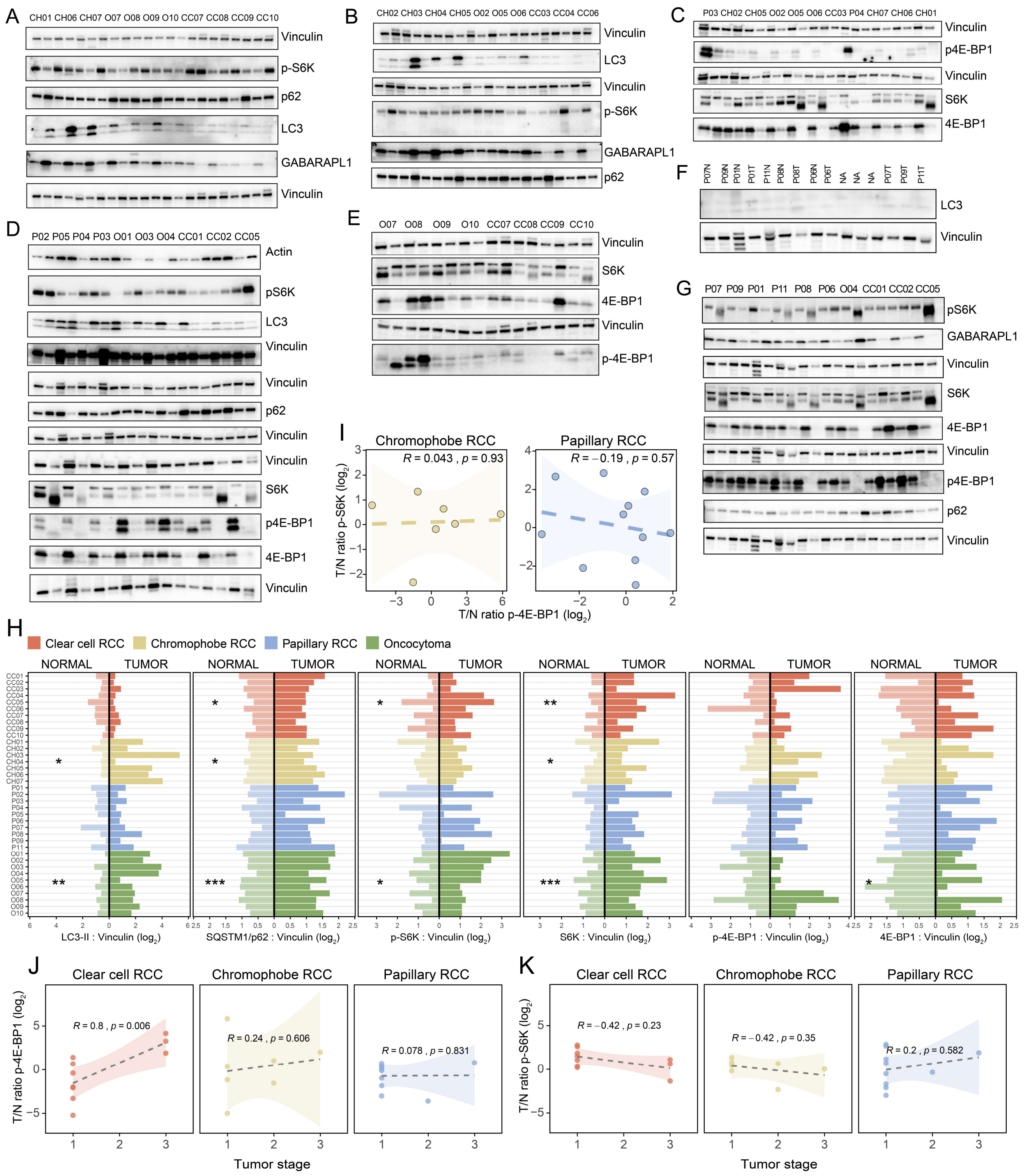


**Supplementary figure 3: A-G**: Raw images of all the western blots that we have used in our analysis. Each subpanel represents one experiment. Several SDS-PAGE gels were run for each experiment, thus one vinculin loading control blot is included for each set of sample loading. Total and phosphorylated forms of 4E-BP1 and S6K were immunoblotted on different membranes. **H**: Quantification of protein levels normalized to vinculin as measured by western blot. Values are $log_2$ transformed. **** = p <0,0001, *** = p <0,001, ** = p < 0,01, * = p <0,05. Non-significant statistical comparisons are not shown. **I**: Relative tumor p-S6K versus relative tumor p-4E-BP1 levels for chromophobe (yellow) and papillary (blue) RCC samples as determined by immunoblotting. Statistical test is linear regression with Pearson's coefficient. Ribbon represents standard error of estimate. **J-K**: Relative tumor level of p-4E-BP1 (J) and p-S6K (K) across tumor stage for RCC subtypes. Values are $log_2$ transformed. Statistical test is linear regression with Pearson's coefficient. Ribbon represents standard error of estimate.

A

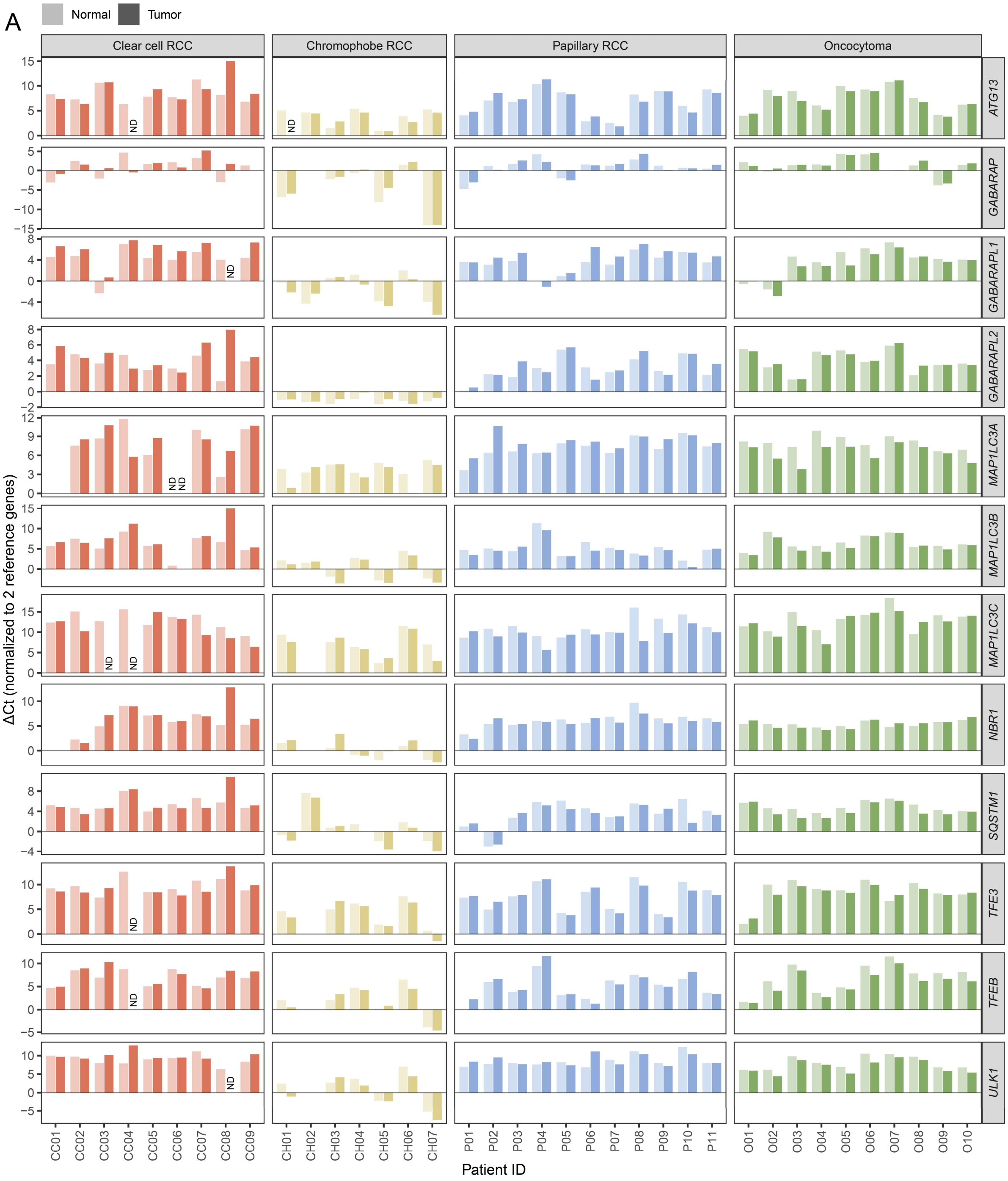


B

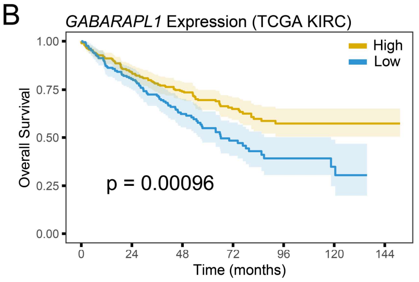


C

| Normalized Mutual Information (k=4) | |
|---|---|
| **Metric used for clustering** | **NMI score** |
| WB (Fig 3E) | 0.365 |
| qPCR (Fig 4A) | 0.280 |
| WB + qPCR + IF (Fig 4H) | 0.575 |

**Supplementary figure 4: A**: RT-qPCR ΔCt values for all patient samples and targets. ND = non-detectable mRNA levels. CC01 is missing values for *NBR1* and *MAP1LC3A*. CH02 is missing values for *ULK1*, *TFEB*, *TFE3*, *NBR1* and *GABARAP*. **B:** Overall survival curves for high and low *GABARAPL1* expression levels in patient samples extracted from KIRC TCGA dataset. Expression cut-off was set to the median expression. **C:** Normalized Mutual Information (NMI) values comparing clustering of subtypes in presented heatmaps.

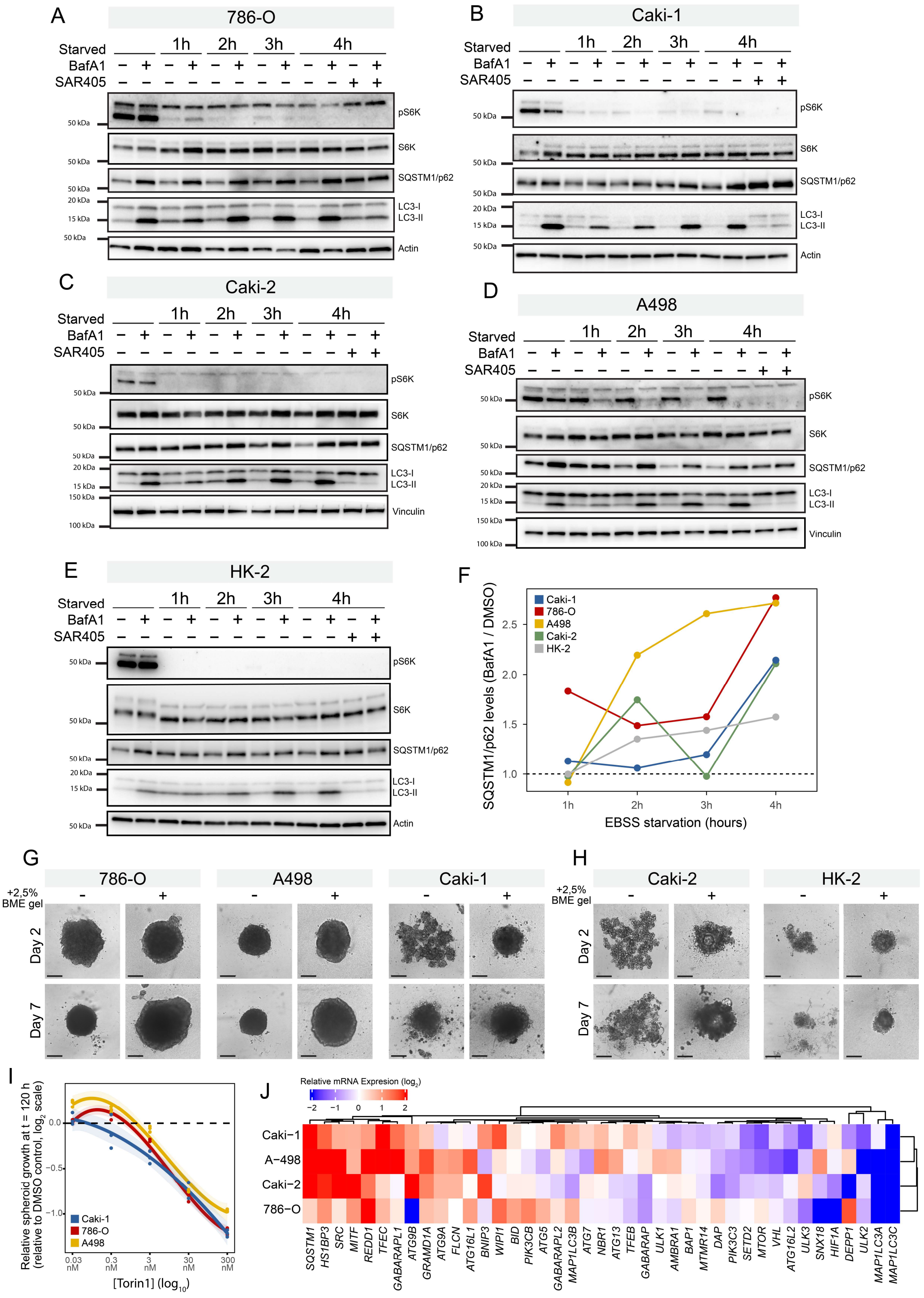


**Supplementary figure 5: A-E**: Western blots to determine autophagy status of several RCC cell lines and kidney proximal tubule cell line HK-2 as control. The cells were starved from 1 to 4 hours and treated with either DMSO, BafA1, SAR405 or a combination thereof. **F**: SQSTM1/p62 levels as measured by immunoblotting in A-E. Values are $BafA_1$/DMSO ratios per timepoint throughout EBSS starvation. Horizontal dotted line represents a ratio value of 1. **G-H**: Formation of spheroids from RCC cell lines and HK-2 with and without addition of 2,5% Basement Membrane Extract (BME) gel. Days are relative to cell seeding at day 0. **I**: Spheroid growth response at t = 120 hours shown as relative growth compared to DMSO control for RCC cell lines Caki-1 (blue), 786-O (red) and A498 (yellow) after drug treatment with Torin1. Confidence bands represent 95% confidence interval. Each datapoint represents a technical replicate. **J**: Heatmap showing relative expression (extracted from CCLE database) of selected genes known to be involved in or important for autophagy. Expression levels are relative to HEKTE expression, the only normal kidney cell line present in CCLE dataset. Relative expression level is shown in $log_2$ scale.
